# Synergistic TOR and ERK inhibition mitigates the hereditary haemorrhagic telangiectasia-like phenotype and excess *kugel* formation in endoglin mutant zebrafish

**DOI:** 10.1101/2021.06.16.448717

**Authors:** Ryan O. Snodgrass, Helen M. Arthur, Timothy J.A. Chico

## Abstract

**Rationale:** Hereditary haemorrhagic telangiectasia (HHT) is an inherited bleeding disorder characterised by arteriovenous malformations (AVMs). Such AVMs affect lungs, liver and brain, whilst telangiectases in mucocutaneous tissues are prone to haemorrhage. HHT type I is caused by loss-of-function endoglin (*ENG*) mutations. Evidence suggests AVMs result from abnormal responses to VEGF signalling.

**Objective:** We therefore characterised the vascular abnormalities in *eng* mutant zebrafish and investigated whether these are prevented by inhibiting different pathways downstream of VEGF signalling.

**Methods and Results:** We used light sheet fluorescence microscopy to visualise the vasculature in *eng*^mu130^ mutant zebrafish. In addition to previously described significantly enlarged dorsal aorta and posterior cardinal vein at 3d post fertilisation, *eng*^mu130^ embryos had an enlarged basilar artery (BA), and increased formation of endothelial *“kugeln”* on cerebral vessels. Adult *eng*^mu130^ fish developed skin AVMs, retinal vascular abnormalities, and an enlarged heart. Tivozanib (AV951), a VEGF receptor tyrosine kinase inhibitor, prevented development of the abnormally enlarged major vessels and normalised the number of *kugeln* in *eng*^mu130^ embryos. Inhibiting discrete signalling pathways downstream of VEGFR2 in *eng*^mu130^ embryos gave further insights. Inhibiting TOR or MEK prevented the abnormal trunk and cerebral vasculature phenotype, whilst targeting NOS and MAPK had no effect. Combining subtherapeutic TOR and MEK inhibition prevented the vascular phenotype, suggesting synergy between TOR and MEK/ERK signalling pathways.

**Conclusions:** These results indicate the HHT-like phenotype in zebrafish endoglin mutants can be mitigated through modulation of VEGF signalling, and implicate combination low dose ERK and TOR pathway inhibitors as a therapeutic strategy in HHT.

**Graphical Abstract:** 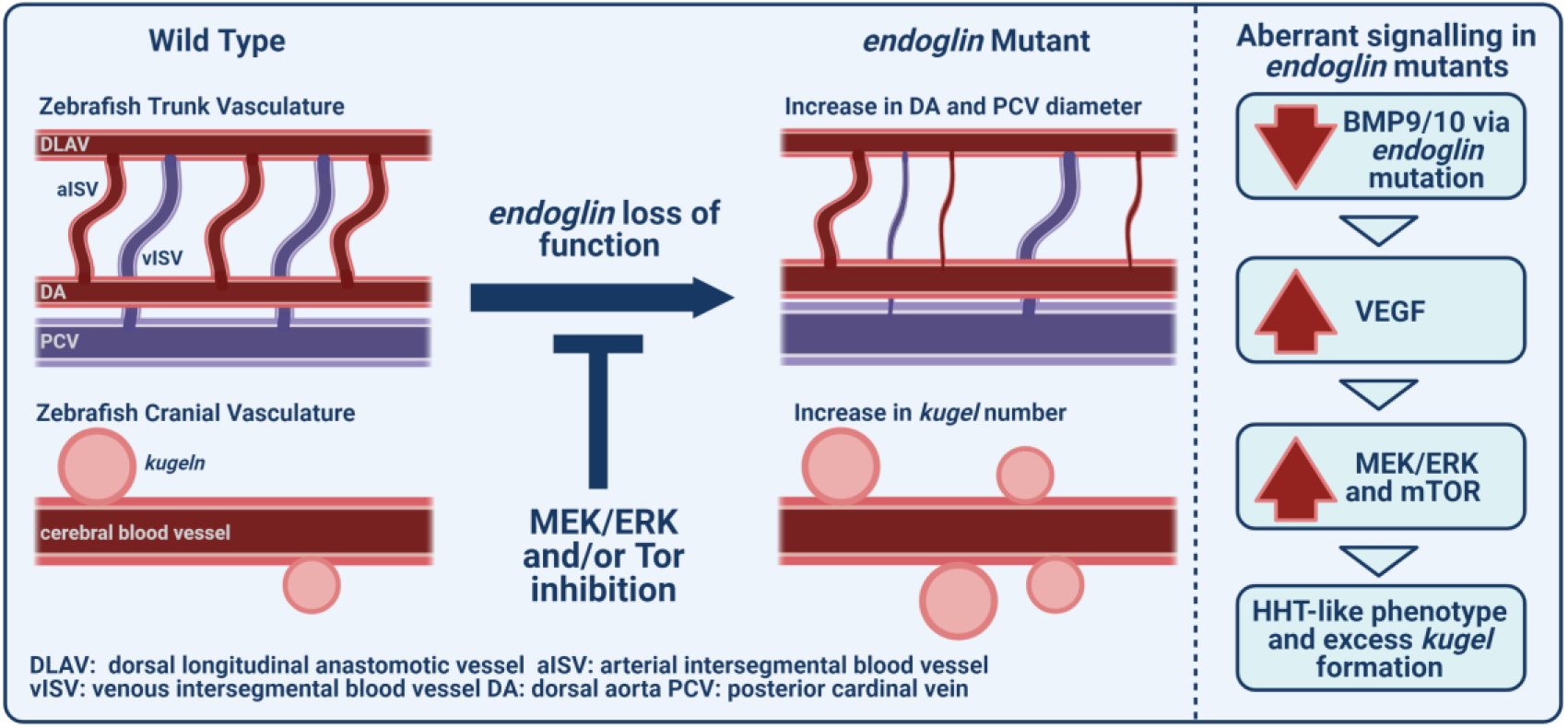

## Introduction

Endoglin (ENG) is highly expressed in endothelial cells (ECs), where it regulates development and maintenance of blood vessels [1–3]. Endoglin is a high affinity co-receptor for BMP9 and BMP10 ligands of the TGF-β superfamily [4, 5]. It behaves as a BMP9/10 ligand reservoir on the EC surface [6] and promotes ligand-induced ALK1 phosphorylation of SMAD1/5/8 [7]. Once phosphorylated, these SMAD proteins can bind to SMAD4 to move to the nucleus where they regulate expression of genes regulating angiogenesis and vascular morphology. Reduced ENG or ALK1 activity due to heterozygous loss of function mutations leads to the human disease hereditary haemorrhagic telangiectasia (HHT) [8, 9]. HHT is characterised by arteriovenous malformations (AVMs), which are local direct connections between arteries and veins that bypass the capillary system, and thereby disrupt blood flow and oxygen circulation. Large AVMs may affect the lungs, liver and brain, whilst telangiectases arise from smaller AVMs between a postcapillary venule and arteriole in mucocutaneous tissues, and are prone to rupture and haemorrhage [10, 11].

Genetic mouse models have been developed to study HHT [12]. Endoglin-null mice die by gestational day 11.5 due to defective blood vessel and heart development [1, 13]. Loss of endothelial ENG in neonatal mice leads to retinal AVMs [2] that result from an abnormal increase in endothelial proliferation and reduced EC migration against blood flow during angiogenesis [2, 3]. A comparable AVM phenotype is seen in neonatal retinas of mice that have lost endothelial Acvrl1, and results from uncoupling of haemodynamic forces to EC migration [14]. If loss of endothelial Eng is delayed until adulthood, this results in peripheral AVMs associated with increased vessel growth leading to high output heart failure. Furthermore AVMs caused by endothelial loss of ENG can be corrected by over-expression of ALK1 [15] confirming the consensus view that ENG is the upstream co-receptor for BMP9/10 in the ENG/ALK1/SMAD1,5,8 signalling pathway.

In zebrafish embryos, loss of *eng* leads to arteriovenous shunting between an enlarged dorsal aorta (DA) and an enlarged posterior cardinal vein (PCV) [16]. These vessels show increased EC size and increased blood flow, with a corresponding reduced flow through the smaller intersegmental vessels [16]. These defects reflect a reduced sensitivity to haemodynamic cues following loss of endoglin [17]. Thus, endoglin mediates multiple EC responses, including proliferation, migration against flow and cellular footprint in the context of angiogenic cues, such as VEGF. As ENG promotes BMP9/10 signalling through ACVRL1, further insights into the importance of this pathway in vascular development and remodelling have come from analysis of *acvrl1* and BMP10 zebrafish mutants. Loss of acvrl1 function leads to increased basilar artery diameter and AVMs in the embryonic cranial vasculature resulting from an accumulation of ECs resulting from reduced migration of ECs towards the heart against blood flow [18]. BMP10 mutant zebrafish embryos also develop enlarged cranial vessels and subsequently develop high cardiac output associated with dermal and hepatic vascular defects [19]. Thus ENG promotes BMP9/10/ACVRL1 signalling to couple haemodynamic cues to enable EC migration against blood flow during development and remodelling of the vasculature. Disruption of this pathway leads to AVMs, and depending on their size and location can sufficiently alter the circulatory system to disrupt normal heart function. However, the underlying molecular mechanisms leading to AVM formation in HHT are still not fully understood, and advances in understanding are required to improve treatment.

Targeting VEGF signalling ameliorates the morbidity of HHT disease. The anti-VEGF inhibitor bevacizumab reduces epistaxis and high cardiac output in HHT patients with hepatic AVMs [20]. Furthermore, blocking VEGF signalling reduces vascular malformations in mouse models of HHT1 and HHT2 [21, 22]. However, recent studies have highlighted the increased risk of serious adverse cardiovascular complications when targeting VEGF signalling in cancer patients [23]. VEGF signalling is complex and drives numerous downstream pathways **[Supplementary Figure 1]**, making it possible that targeting specific downstream pathways may provide benefit with reduced risk of side-effects. To address which pathway (or combination of pathways) would efficiently achieve this goal, we took advantage of the zebrafish embryo’s suitability for drug-screening assays, including its rapid cardiovascular development, ease of drug administration and transgenic tools for imaging. The *Tg(kdrl:Hsa.HRAS-mCherry)^s916^* transgenic background labels the EC membrane, facilitating live imaging of the embryonic trunk and cranial vasculature, and imaging of the adult retinal vasculature in explanted tissue. We recently used this line to describe a novel EC behaviour, in which the cerebral vessels form structures termed *kugeln*. *Kugeln* are transient bleb-like structures that extrude abluminally from cerebral blood vessels in the absence of blood-flow, and contain no cytoplasm [24]. They have only been detected in cerebral vessels, and although their function is currently unclear, they may provide new insights into mechanisms of cerebrovascular development. Here, in addition to previously reported enlargement of major trunk vessels in *eng* mutant zebrafish embryos, we discovered an increase in *kugel* formation as well as increased basilar artery diameter. We pharmacologically inhibited either global VEGF signalling or components of different pathways downstream of VEGFR2 in zebrafish *eng* mutant and control embryos **[Supplementary Figure 1]** and identified MEK/ERK and mTOR as synergistic therapeutic targets in HHT.

## Methods

### Zebrafish Husbandry

Animal experiments were performed at the University of Sheffield under Home Office project licence PPL 70/8588. The *endoglin* mutant (*eng*^mu130^) [16] was kindly provided by Dr Arndt Siekmann. *eng*^mu130^ lines were raised in *Tg(kdrl:Hsa.HRAS-mCherry)^s916^* [25] and *Tg*(*kdrl:EGFP*)^y7^ [26] transgenic backgrounds, which label the EC membrane and endothelial cytoplasm respectively. Adult fish were housed in a recirculating aquarium with a 14-hour light / 10-hour dark cycle at 28.0±1°C, pH 7.5, oxygen saturation 80%. Clutches of sibling embryos were generated by pair-mating male and female heterozygous adults to generate mixed wild-type (WT), heterozygous (+/-), and homozygous (-/-) mutant offspring.

### Morphology of adult zebrafish hearts and retinae

*Tg(kdrl:Hsa.HRAS-mCherry)^s916^* adult fish (5-6 months) were killed by a lethal dose of tricaine (MS-222). Freshly dissected hearts were fixed in 4% paraformaldehyde overnight at 4°C. Hearts were weighed, and imaged using a stereomicroscope (Olympus IX81). To prepare retinae, heads were fixed in 4% paraformaldehyde overnight at 4°C. Eyes were enucleated and retinae dissected, flat-mounted onto glass slides and imaged using a fluorescent stereomicroscope (Axio zoom V.16). Number of capillary interconnections, optic artery diameter and vessel branching was quantified as previously described [27].

### Genotyping zebrafish embryos

After experimental observations were complete, anaesthetised embryos were individually placed in microcentrifuge tubes with 25μl of 50mM NaOH, heated to 95°C for 10 minutes, then cooled to 10°C. The reaction was neutralised by addition of 0.5μl 100mM Tris-HCl pH9.5 (Sigma-Aldrich). Genomic DNA was amplified by PCR (MultiGene™ OptiMax Thermal Cycler) as previously described [16] using primers [FWD: GCTGATTAGGGCTGCAAGA, REV: TGTTGTGGTAATTTTACAGTTGCT] to generate a 418bp DNA fragment. Restriction digest at 37°C for 20 minutes with *Msp*1 (New England Biolabs) was performed to cleave the wildtype, but not *eng*^mu130^, PCR product into 246bp + 172bp fragments, and visualised using agarose gel electrophoresis.

### Quantifying embryonic blood vessel diameter

Zebrafish embryos aged 2-4 days post fertilisation (dpf) were imaged for red and/or green fluorescence using a fluorescent stereomicroscope (Zeiss Axiozoom V.16) then anaesthetised using 0.4% tricaine, and mounted in 1% low melting point agarose (LMP) (Biolabs) within a 1mm diameter glass capillary. Samples were suspended in the chamber of a Zeiss Z1 lightsheet microscope filled with E3 medium at 28°C. Samples were excited with a 488nm and 561nm wavelength lightsheet, and the emitted GFP and RFP signals detected using an LP560 filter. Image acquisition and processing was performed using ZEN Black software (Zeiss). Z-stacks of the trunk and head vasculature were used to generate maximum intensity projections (MIPs). Diameters of the dorsal aorta (DA) and posterior cardinal vein (PCV) were measured using Fiji image analysis software [28] at the midway point between 2 intersegmental blood vessels (ISVs) along the yolk extension, five points in total, to generate mean diameter per embryo as previously described [16]. For ISV diameter, three measurements were made along three ISVs between the DA and dorsal longitudinal anastomotic vessel (DLAV) to calculate average ISV diameter for both arterial and venous ISVs. Diameter of the basilar artery (BA) was measured at three points along the vessel, and mean diameter calculated for each embryo. *kugeln* were identified and counted as previously described [24].

### Heart rate measurement

Individual non-anaesthetised embryos were observed under a bright field stereomicroscope (Olympus IX81). Heart rate was calculated over 15 seconds, three times per embryo, expressed as beats per minute (bpm).

### Chemical treatments

All chemicals were dissolved in E3 medium and administered by immersion from 24 hours post fertilisation (hpf) to 48hpf. VEGF signalling was inhibited using 25nM Tivozanib (AV-951, AVEO pharmaceuticals), vehicle control groups were exposed to 0.0025% DMSO. TOR signalling was inhibited using 2-2.5μM Rapamycin (Sigma-Aldrich), vehicle control groups were exposed to 0.2-1% DMSO. MEK signalling was inhibited using 7.5-10μM PD0325901 (Sigma-Aldrich), vehicle control groups were exposed to 0.75-1% DMSO. p38 MAPK signalling was inhibited by 25μM SB 203580 (Sigma-Aldrich). Nitric oxide synthase (NOS) inhibition was achieved by incubation with 0.5mM L-NAME (Sigma-Aldrich) diluted in E3 medium.

### Statistical analysis

Statistical analysis was performed in GraphPad Prism 7. Data were subjected to D’Agostino-Pearson normality test before analysis. Statistical tests used are indicated in figure legends. Error bars display standard deviation (S.D.). Each experiment was repeated three times, unless otherwise stated. P values are indicated as follows: *=<0.05, **=<0.01, ***=<0.001, ****=<0.0001.

## Results

### *endoglin* mutant zebrafish display cardiovascular defects

Although *eng*^mu130^ embryos are similar in size to their wild type siblings **[Supplementary Figure 4]**, adult *eng*^mu130^ mutant zebrafish adults appear smaller and weighed less than WT siblings [**Figure 1 A,C**]. However, hearts were significantly larger in *eng* mutants [**Figure 1 B,D**], similar to the phenotype observed in endothelial-specific *Eng* knockout adult mice [21]. Given the prevalence of skin vascular abnormalities in HHT patients, we examined *eng* mutants for similar cutaneous phenotypes. 71% (5/7) of adult *eng*^mu130^ zebrafish displayed cutaneous blood vessel abnormalities, which were absent in WT (0/5) [**Figure 1 A**]. We observed extensive retinal vessel malformations in adult *eng*^mu130^ mutant zebrafish [**Figure 1 E,G,H,I**], including increased capillary interconnections, increased optic artery diameter and increased vessel branching compared with WT.

**Figure 1.**
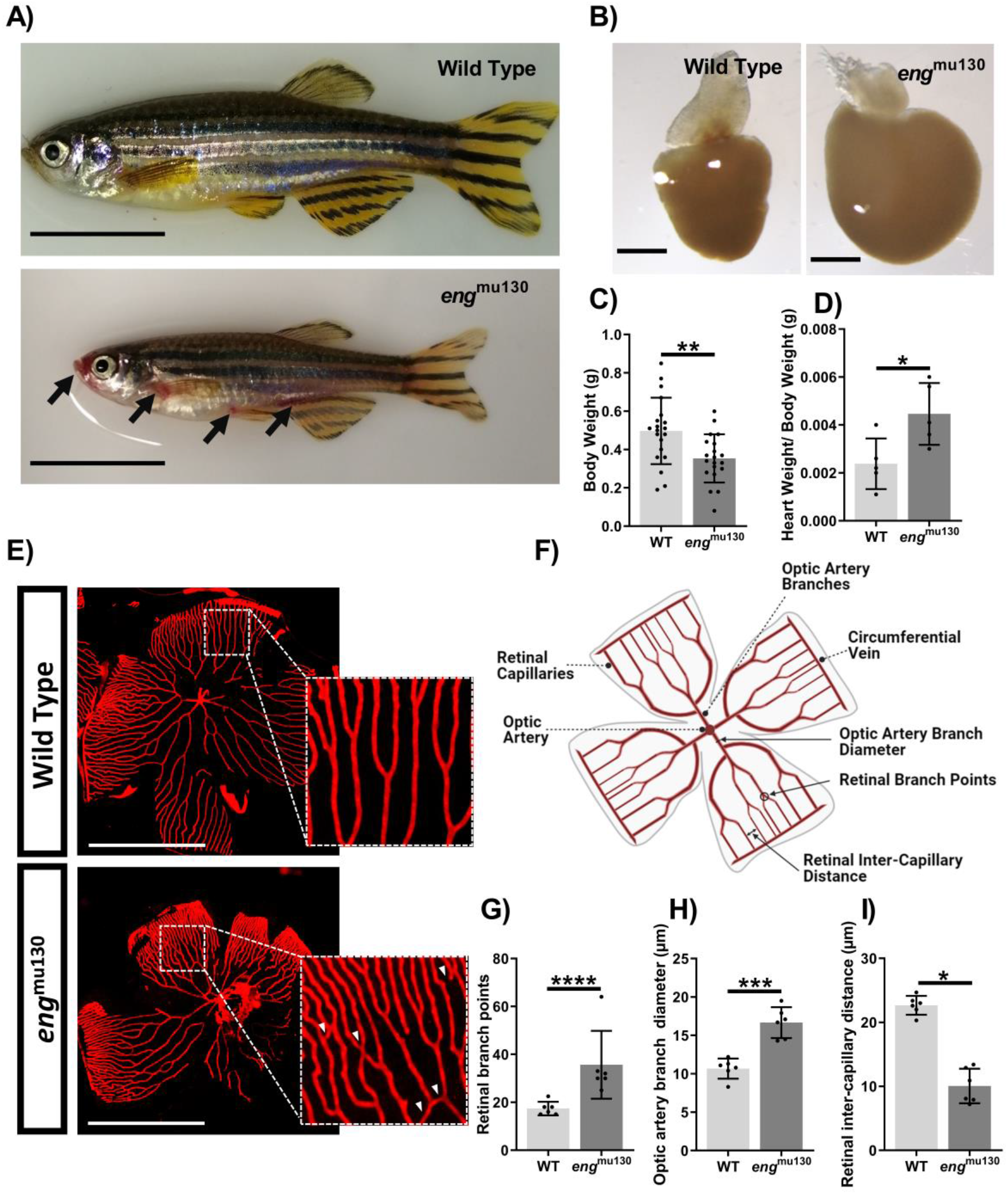
Adult *eng*^mu130^ fish display cutaneous and retinal vascular malformations similar to HHT and an enlarged heart. A) Adult 5-6 month old *eng*^mu130^ zebrafish display cutaneous vascular malformations (black arrows). Scale bar = 1cm. B) Representative *eng*^mu130^ and wild type hearts (n=5/group). Scale bar = 500μm. C) Body weight of adult WT and *eng*^mu130^ zebrafish D) Heart weight/body weight of WT and *eng*^mu130^ zebrafish. E) Fluorescent micrographs of whole mount retinae from *Tg*(*kdrl:Hsa.HRAS-mCherry*)^s916^ zebrafish showing abnormal vessel communications in *eng*^mu130^ (white arrowheads). F) Schematic diagram of a 5-6 month old zebrafish retinal vasculature. Figure created with BioRender (https://biorender.com/). G) Number of retinal vascular branches in WT and *eng*^mu130^ homozygous siblings. H) Optic artery diameter in WT and *eng*^mu130^ homozygous siblings. I) Inter-capillary distance in WT and *eng*^mu130^ homozygous siblings. (Unpaired Student’s t-test, 5-7 animals/group.)

In normal zebrafish development, the DA and PCV diameter reduces between 52 and 100hpf, a process previously shown to be defective in *eng*^mu130^ mutants [16]. In agreement with this, we found the DA and PCV diameters were significantly larger in *eng*^mu130^ mutants compared with WT, a phenotype most pronounced in the older (100hpf) embryos **[Supplementary Figure 2 C,D]**. This is due to an increase in EC size and not increased EC number [16] **[Supplementary Figure 3]**. Increased flow through the enlarged major vessels in *eng^mu130^* mutants leads to correspondingly reduced flow in ISVs, which leads to a delayed opening of their lumens [16], a finding we also confirm **[Supplementary Figure 2 E,F]**. Thus, *eng* plays an important role in determining vessel calibre in the zebrafish embryo.

To determine whether developmental abnormalities were seen in additional vascular beds, we examined the zebrafish cranial vasculature at 52–100hpf. The basilar artery (BA) had an increased diameter, but no increase in EC number, in *eng*^mu130^ mutants compared with WT siblings **[Figure 2 A,C,D]**. This is similar to the enlarged BA leading to AVM formation in *alk1* and bmp10 mutant zebrafish [18, 19]. The organisation and diameter of the remaining cerebral vessels was very similar between *eng*^mu130^ and WT embryos from 48-96hpf **[Figure 2 B]**. We also observed that *eng*^mu130^ mutants developed significantly more cerebral vessel *kugeln* compared to WT siblings **[Figure 2 E,F]**. Thus, in line with human HHT, there are a range of abnormal vascular phenotypes in zebrafish *endoglin* mutants.

**Figure 2.**
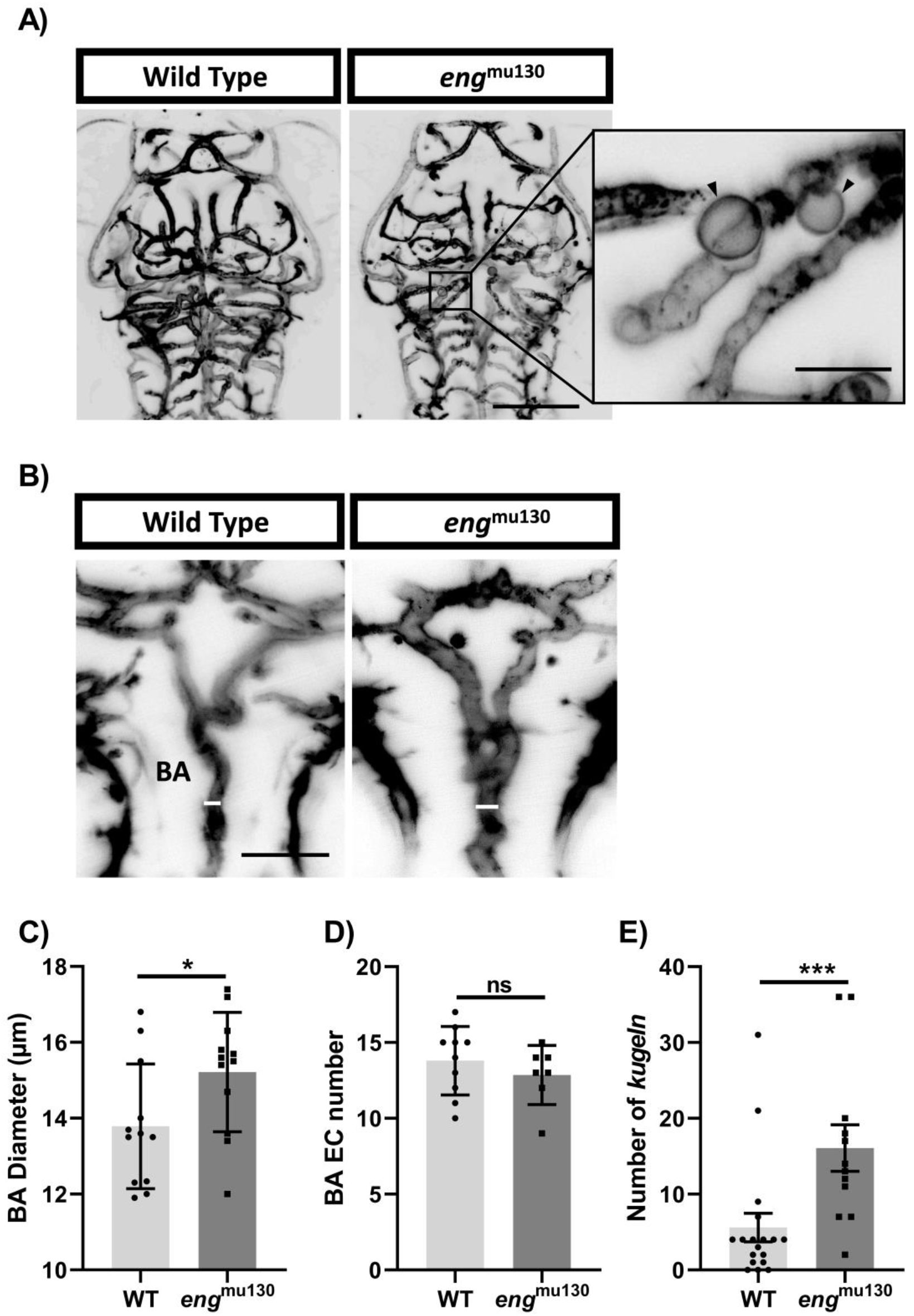
*eng*^mu130^ mutant embryos have increased basilar artery diameter and increased number of endothelial *kugeln*. A) Representative maximum intensity projection of the basilar artery (BA) at 72hpf. Scale bar = 50μm. B) Representative maximum intensity projection of *Tg(kdrl:Hsa.HRAS-mCherry)^s916^* zebrafish brain vasculature at 72hpf. Black arrowheads indicate individual *kugeln*. Scale bar = 150μm/50μm. C) BA diameter in *eng*^mu130^ mutants and WT siblings (unpaired Student’s t-test, 10-12 animals/group). D) *endoglin* mutation did not alter EC number in the BA (unpaired Student’s t-test, 10-12 animals/group). E) *Kugel* number per animal at a single timepoint was increased in *eng*^mu130^ mutants (Mann–Whitney U-test, 12-18/group). F) *Kugel* diameter was not different in *endoglin* mutants (Student’s *t*-test, 12-18/group.)

### Abnormal brain and trunk vasculature of *eng^mu130^* zebrafish embryos is prevented by VEGFR2 inhibition

As loss of endoglin disrupts integration of BMP9/10 and VEGF signalling pathways, we investigated VEGF signalling in *eng*^mu130^ zebrafish in more detail. First, treating *eng*^mu130^ embryos between 2dpf and 3dpf with the VEGFR2 receptor tyrosine kinase inhibitor AV951 (Tivozanib) prevented progression of the DA and PCV mutant phenotypes, but did not affect the vessel calibre of WT animals **[Figure 3 B,C,D]**. This indicates the HHT-like phenotype in zebrafish *eng* mutants can be mitigated by inhibiting VEGF signalling. We next asked whether the abnormal cerebral phenotype of *endoglin* mutants is caused by a similar dysregulation of VEGF signalling. Treating *eng*^mu130^ embryos between 2dpf and 3dpf with AV951 prevented increased *kugel* formation in the mutant embryos, but did not affect *kugel* number in WT siblings **[Figure 3 G]**. The increased BA diameter in endoglin mutants was also prevented by treatment with AV951 **[Figure 3 H]**.

**Figure 3.**
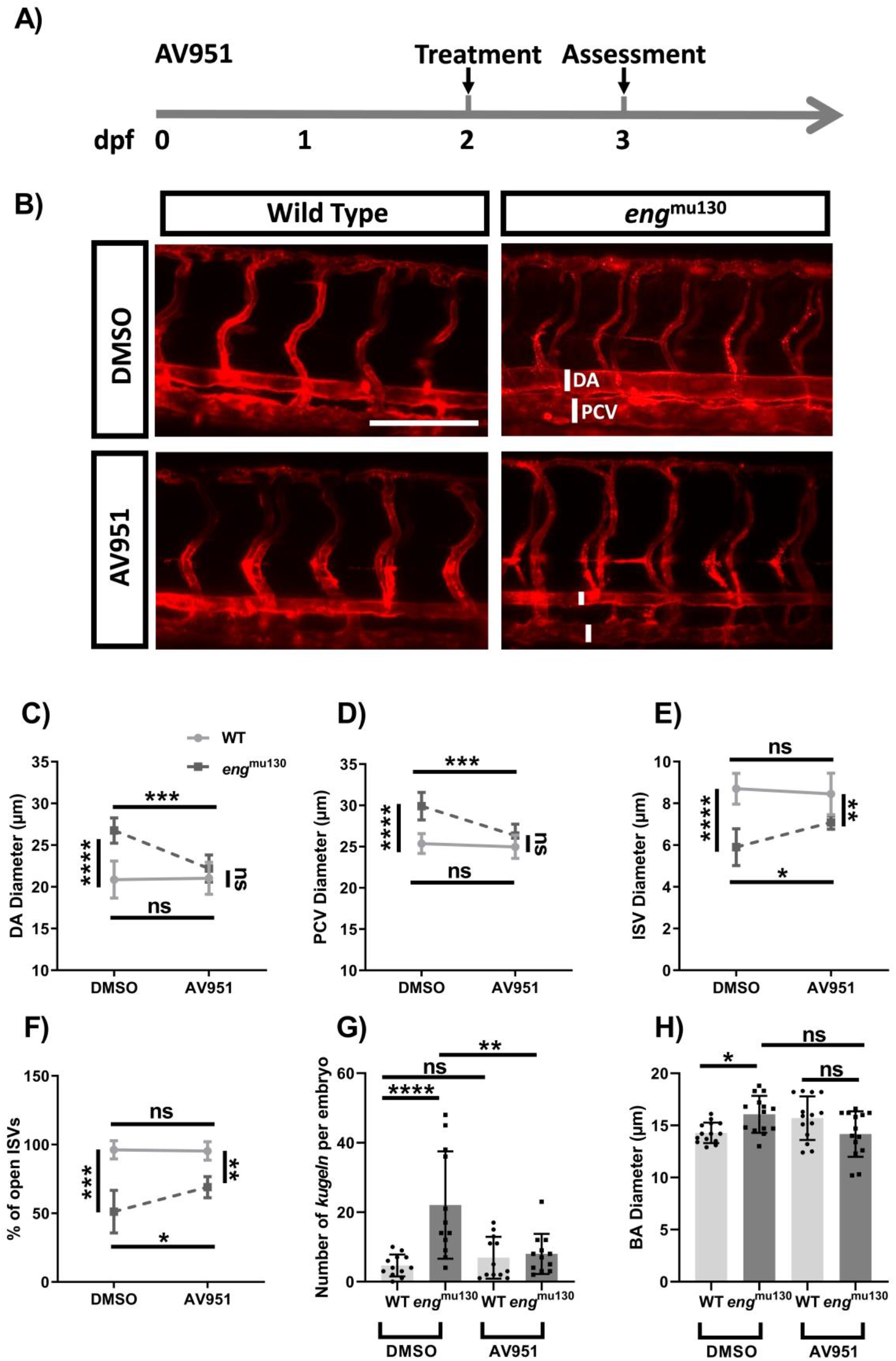
VEGF inhibition between 2-3dpf rescues the abnormal trunk and cerebral vessel phenotypes of *eng*^mu130^ mutants. A) Experimental plan for rescuing *endoglin* mutant phenotype in zebrafish using AV951 (dpf, days post fertilization) B) Representative maximum intensity projection of *eng*^mu130^ and WT embryos ± AV951 (25nM) treatment for 24h. Scale bar = 150μm. C) DA diameter in *eng*^mu130^ and WT embryos ± AV951 treatment. D) PCV diameter in *eng*^mu130^ and WT embryos with and without AV951 treatment. E) ISV diameters in *eng*^mu130^ and WT embryos ± AV951 treatment. F) Percentage of open ISVs in *eng*^mu130^ and WT embryos ± AV951 treatment. G) Number of *kugeln* in *eng*^mu130^ and WT embryos ± 25nM AV951 treatment. H) BA diameter in *eng*^mu130^ and WT embryos ± 25nM AV951 treatment. (two-way ANOVA with Tukey post-hoc test, 10/group.)

### TOR and MEK inhibition prevents abnormal brain and trunk vasculature of *eng^mu130^* zebrafish embryos

VEGFR2 signalling is complex and signals downstream via several pathways. In vitro, BMP-9 decreases VEGF signalling, and loss of BMP9 signalling through ACVRL1 or ENG depletion leads to increased pERK levels[21, 29]. Treatment of embryos at 2dpf for 24h with 10μM PD0325901 to inhibit MEK, upstream of ERK, prevented the enlarged DA and PCV of *eng*^mu130^ embryos without affecting vessel diameter of WT siblings **[Figure 4 A,B]** Furthermore, 10μM PD0325901 prevented the abnormally increased number of *kugeln* in *eng*^mu130^ embryos, but did not affect *kugel* number in WT animals. This suggests that upregulation of MEK/ERK signalling due to *eng* loss of function contributes to the trunk vessel abnormalities and increased *kugel* formation in *eng* mutants **[Figure 4 E]**.

**Figure 4.**
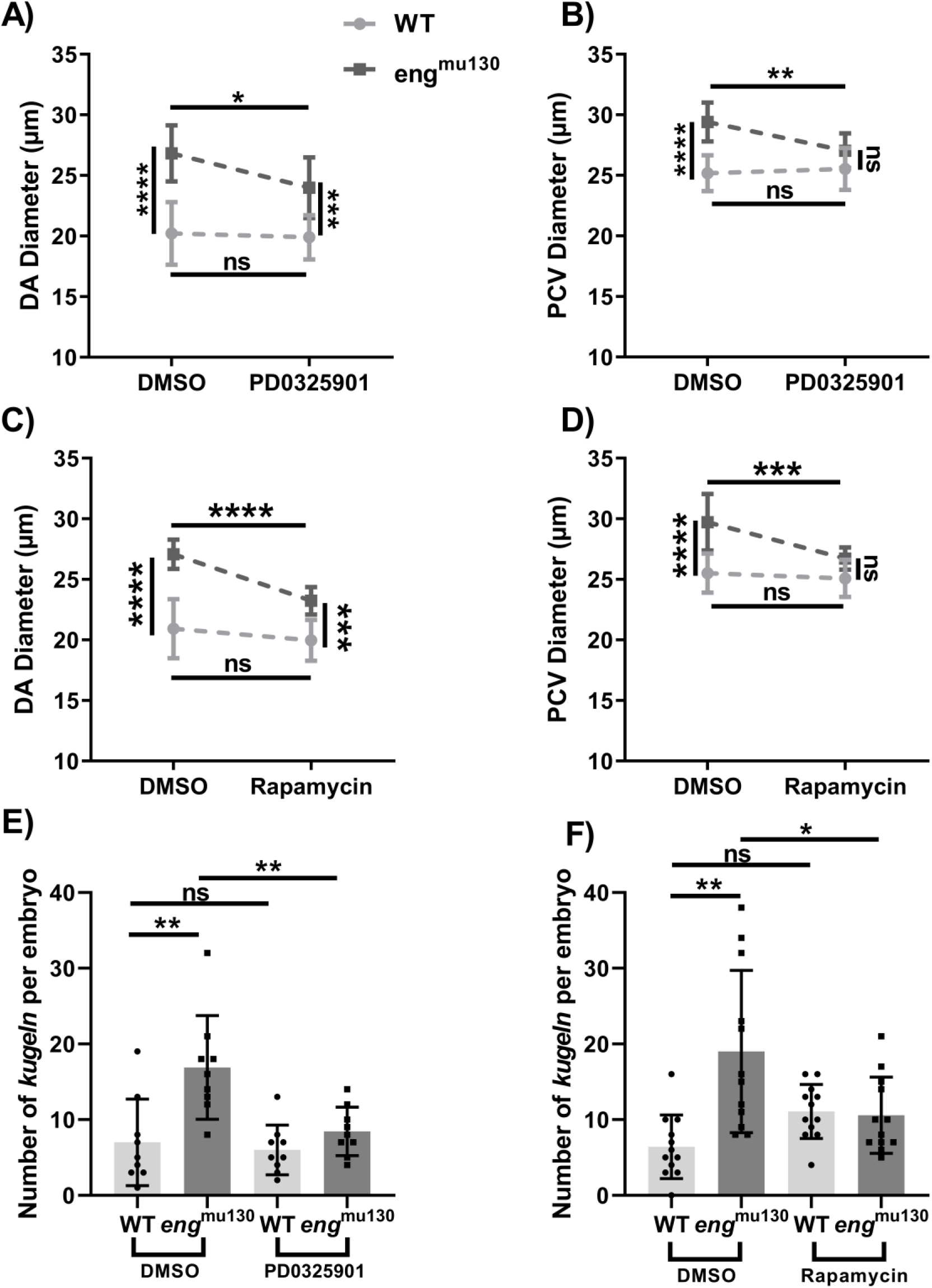
Either TOR or MEK inhibition normalises the vascular phenotype of *eng*^mu130^. WT and *eng*^mu130^ mutant embryos were treated with the TOR inhibitor Rapamycin (2.5μM) or with the MEK inhibitor PD0325901 (10μM) for 24h between 2-3dpf, or with DMSO vehicle. A/B) *eng*^mu130^ mutant embryos show reduced DA and normal PVC diameter following PD0325901 treatment. D/E) *eng*^mu130^ mutant embryos show reduced DA diameter and normal PCV diameter following rapamycin treatment. E) *eng*^mu130^ mutant embryos show normalised *kugel* formation following PD0325901 treatment. F) *eng*^mu130^ mutant embryos show reduced *kugel* formation following Rapamycin treatment. (two-way ANOVA with Tukey post-hoc test. n = 10-14/group.)

A second major pathway downstream of VEGFR2 is the PI3K/AKT pathway, which affects both mTOR and eNOS function to regulate cell survival and vasoregulation, respectively **[Supplementary Figure 1]**. Targeting PI3K can reduce AVMs in murine HHT2 models [17]. Furthermore, in a model of HHT caused by loss of BMP9 and BMP10 ligands, endothelial mTOR levels are increased and targeting mTOR reduces vascular defects [30]. Therefore, we assessed the effect of rapamycin (also known as Sirolimus) in inhibiting TOR in *eng*^mu130^ zebrafish embryos and found that both DA and PCV diameters were normalised whilst WT embryos were unaffected **[Figure 4 C,D]**. Rapamycin also prevented the increase in *kugel* number in *endoglin* mutant embryos **[Figure 4 F]**. Endothelial NOS is a critical enzyme regulating the levels of nitric oxide (NO) available for diffusion to neighbouring smooth muscle cells (SMCs) where it leads to vasodilation. eNOS is upregulated both by flow and by VEGF signalling, and there is evidence that endoglin regulates coupling of eNOS activity [31]. However, we found no detectable effect on DA or PCV vessel size in *eng*^mu130^ embryos following treatment with L-NAME, consistent with the lack of vascular smooth muscle cells supporting these vessels at this stage of development. **[Supplementary Figure 5]**.

Finally, we investigated if combined low dose mTOR and MEK inhibition prevents abnormal trunk vasculature in *eng*^mu130^ embryos. Neither 2μM Rapamycin nor 7.5μM PD0325901 alone altered the vascular phenotype of *eng*^mu130^ mutants, but co-treatment normalised both DA and PCV diameters and reduced the excess kugel formation phenotype **[Figure 5 A,B, C]**, suggesting that upregulation of both MEK/ERK and AKT/mTOR pathways contribute to the abnormal vessel phenotype in eng mutants.

**Figure 5.**
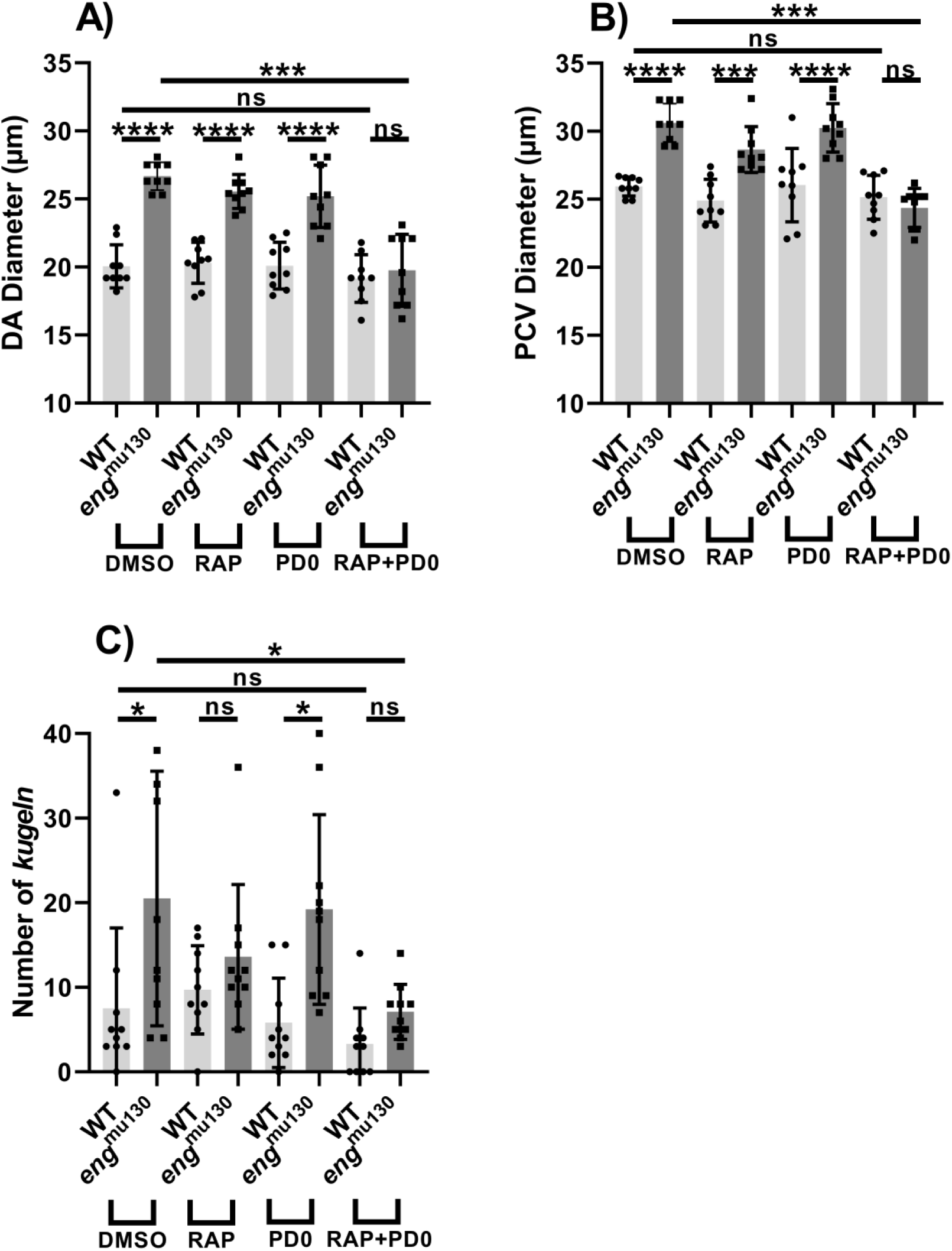
Combined low dose TOR and MEK inhibition prevents abnormal trunk vasculature and excess *kugel* formation in *eng*^mu130^ zebrafish embryos. A,B, C) Neither 2μM Rapamycin nor 7.5μM PD0325901 alter the vascular phenotype of *eng*^mu130^ mutants, but combined treatment resulted in normalised DA and PCV diameters, as well as reduced kugel formation in *eng*^mu130^ zebrafish embryos. (two-way ANOVA with Tukey post-hoc test (10-14/group.)

## Discussion

Our findings substantially expand the previously described vascular phenotype in *eng*^mu130^ mutant zebrafish. Adult *eng* mutants display cutaneous vascular lesions similar to HHT patients, retinal vascular abnormalities and cardiac enlargement. In embryos we uncover a striking cerebrovascular phenotype of enlarged basilar artery (BA) and increased endothelial *kugel* formation in *eng* mutants. *Kugeln* are large transient spherical structures protruding from cerebral blood vessels that are Notch-dependent, NO-enriched, and VEGF-sensitive [24]. Currently, the physiological significance of *kugeln* is not clear, but this is the first work that implicates *kugel* formation in a model of human disease.

Treatment of *eng*^mu130^ embryos with Tivozanib to target VEGFR2 prevented development of the abnormally enlarged major vessels and normalised cerebrovascular *kugel* number. This indicates that the HHT-like phenotype in zebrafish *eng* mutants can be mitigated by inhibiting VEGF signalling, in agreement with other studies in mouse models of HHT [3, 21, 22] and current clinical therapies targeting VEGF in HHT patients [20]. However, anti-VEGF treatment has to be continuous or semi-continuous to maintain suppression of abnormal angiogenic responses, and anti-VEGF therapies such as Bevacizumab have severe side effects and require repeated intravenous delivery [23]. Furthermore, VEGF signalling activates numerous downstream pathways **[Supplementary Figure 1]** and knowing which of these is dysregulated in HHT will help to guide more focused therapeutic interventions.

We therefore took advantage of the zebrafish embryo’s suitability for drug-screening assays, including its rapid cardiovascular development, *ex utero* fertilisation and rapid uptake of inhibitors from the incubating medium to target separate signalling pathways downstream of VEGFR2 in *eng*^mu130^ embryos. As *Eng* mutants show increased phosphorylated ERK (pERK) activity in endothelial cells downstream of VEGF signalling and increased pERK was observed in ECs associated with AVMs [21] this may represent a key pathway to target downstream of VEGF. Indeed, sporadic human cerebral AVMs also have an exaggerated pERK response suggesting aberrant pERK activation is a common feature of AVMs [32, 33]. Treatment of embryos for 24h between 2 and 3dpf with PD0325901 to inhibit MEK, upstream of ERK, normalised the enlarged vessels and reduced the excess *kugel* formation in *eng*^mu130^ zebrafish embryos. These timings were chosen as pERK is present at an early stage (from 18hpf) in the developing DA, and predefines ECs that are going to contribute to ISV formation [34]. However, this initial process is completed by 2d when MEK inhibition was initiated. Importantly, development and vessel size (including ISV) of wild type embryos was unaffected by exposure to PD0325901. Thus, targeting MEK effectively prevented occurrence of the *eng* mutant vascular phenotype. In contrast, targeting p38MAPK, a second pathway downstream of VEGF signalling, had no detectable effect on the enlarged trunk vessels in *eng* mutants, suggesting that altered p38 regulated cell migration was not involved in the HHT phenotype. Finally, we looked at the PI3K/AKT pathway downstream of VEGF. Inhibition of PI3K in *Acvrl1* mouse neonates has previously been shown to protect against development of vascular abnormalities [29, 35]. *Acvrl1*-deficient endothelial cells show enhanced activation of the PI3K/AKT pathway [15, 29, 35], which would be expected to increase both eNOS and mTOR activity downstream. We therefore next targeted NOS using L-NAME, but found no detectable effect on DA or PCV vessel size in *eng*^mu130^ embryos. Although previous evidence has suggested eNOS may be involved in HHT due to eNOS uncoupling in Eng mutant endothelial cells [31], our data would suggest that increased NOS activity in the developing zebrafish vasculature does not drive vascular malformations, at least not prior to the recruitment of vascular smooth muscle. Finally, we targeted TOR downstream of pI3K/AKT and found that rapamycin treatment normalised the enlarged vessels and reduced the excess *kugel* formation in *eng*^mu130^ embryos. This is consistent with previous findings in a mouse neonatal retinal model caused by loss of BMP9/10 ligands where targeting mTOR had beneficial effects [30]. We therefore show that independently targeting two separate pathways MEK/ERK and mTOR downstream of VEGF could prevent the mutant vascular phenotype in eng^mu130^ embryos. We then asked whether there was synergy between these two pathways by combining low dose TOR and MEK inhibitors. At low dose neither TOR nor MEK inhibitor had any detectable effect on the developing vasculature of *eng* mutant embryos. However, when used together in combination they efficiently reduced the vascular phenotype. These results indicate that the HHT-like phenotype in zebrafish *eng* mutants can be mitigated through modulation of VEGF signalling and implicate synergistic targeting of ERK and mTOR pathways as therapeutic strategy in HHT. The ability to combine subtherapeutic doses of these inhibitors might reduce the risks of toxicity while providing amelioration of the consequences of HHT.

## Non-standard Abbreviations and Acronyms

(HHT): Hereditary haemorrhagic telangiectasia
(AVMs): Arteriovenous malformations
(*ENG*): Endoglin
(BA): Basilar artery
(EC): Endothelial cell
(VEGF): Vascular endothelial growth factor
(DA): Dorsal aorta
(PCV): Posterior cardinal vein
(dpf): days post fertilisation
(hpf): hours post fertilisation
(MIP): Maximum intensity projection
(ISVs): Intersegmental blood vessels
(DLAV): Dorsal longitudinal anastomotic vessel
(BMP): Beats per minute
(NO): Nitric oxide
(SMC): Smooth muscle cell
(pERK): Phosphorylated ERK

## Funding

R.O.S. is funded by an MRC DiMeN studentship; research in T.J.A.C. and H.M.A. laboratories is funded by the British Heart Foundation.

## Data Availability Statement

Data are available upon request.

## Conflicts of Interest

The authors declare no conflict of interest.

## Notes

### Competing Interest Statement

The authors have declared no competing interest.

