## Supplementary data for "Synergistic TOR and ERK inhibition mitigates the hereditary haemorrhagic telangiectasia-like phenotype and excess *kugel* formation in endoglin mutant zebrafish"

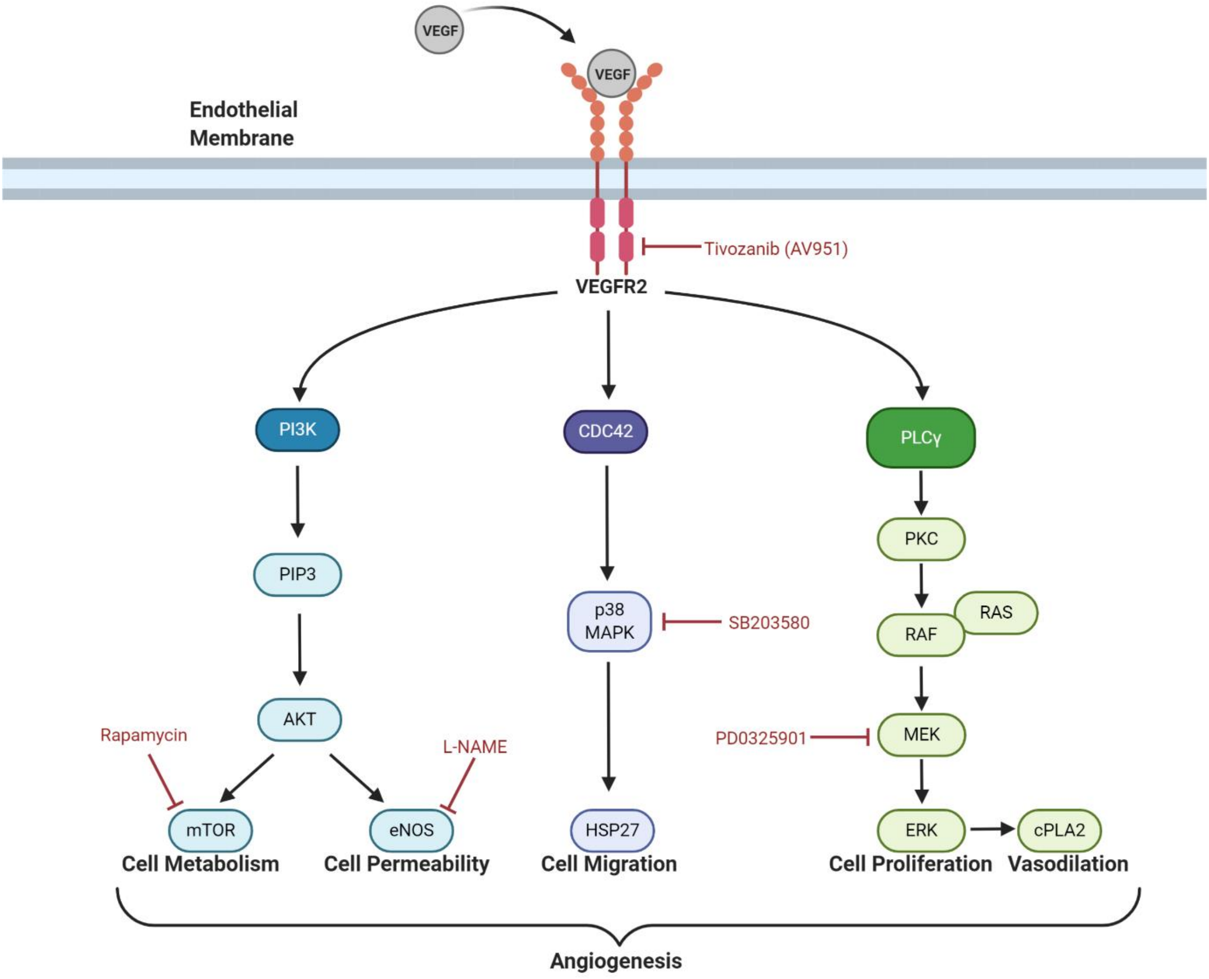


**Supplementary Figure 1 Schematic diagram summarising VEGF signalling in endothelial cells.** Drugs targeting VEGFR2 and enzymes in different downstream pathways are shown in red. Figure created with BioRender (https://biorender.com/). (Adapted from [36]).

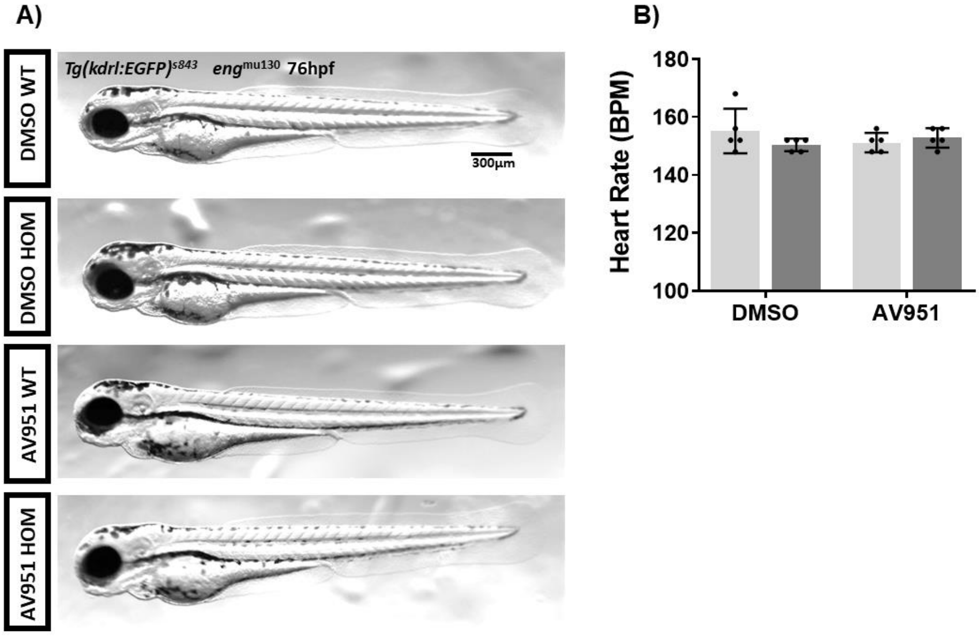


**Supplementary Figure 2 WT and *eng*$^{mu130}$ mutant embryos display no obvious morphological differences when treated with VEGF inhibitors.** A) Representative images of 3dpf WT and *eng*$^{mu130}$ mutant embryos with and without AV951 (50nM) treatment for 24h. B) Quantification of heart rate for WT and *eng*$^{mu130}$ mutant embryos with and without AV951 treatment. n = 5/group.

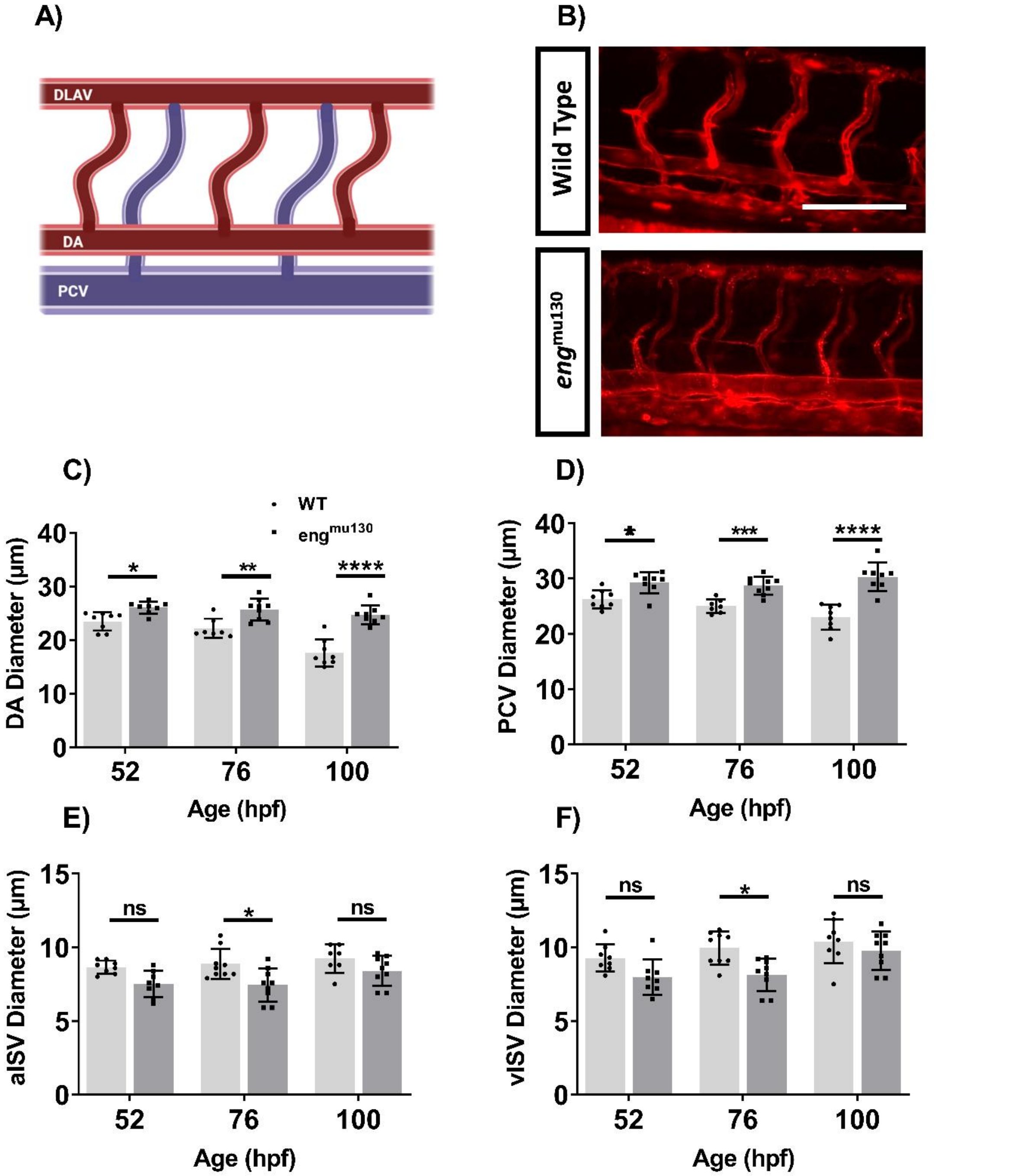


**Supplementary Figure 3 *eng*$^{mu130}$ mutant embryos display increased diameters in the dorsal aorta and cardinal vein compared with control siblings.** A) Schematic diagram of the region of interest. Figure created with BioRender (https://biorender.com/). B) Representative maximum intensity projection of *Tg(kdrl:Hsa.HRAS-mCherry)*$^{s916}$ WT and *eng*$^{mu130}$ trunk vasculature at 76hpf. Abbreviations: DA, dorsal aorta; PCV, posterior cardinal vein; aISV, arterial intersegmental vessel; vISV, venous intersegmental vessel. Scale bar = 100µm. C,D,E and F) Significantly enlarged vessel diameters of DA and PCV in *eng*$^{mu130}$ fish compared with WT siblings. Imaging of eGFP intensity used to establish ISV lumen size. aISV, and vISV are thinner in *eng*$^{mu130}$ mutants at 76h, and partially normalise by 100h. (one-way ANOVA with Tukey post-hoc test. n = 8-10/group.)

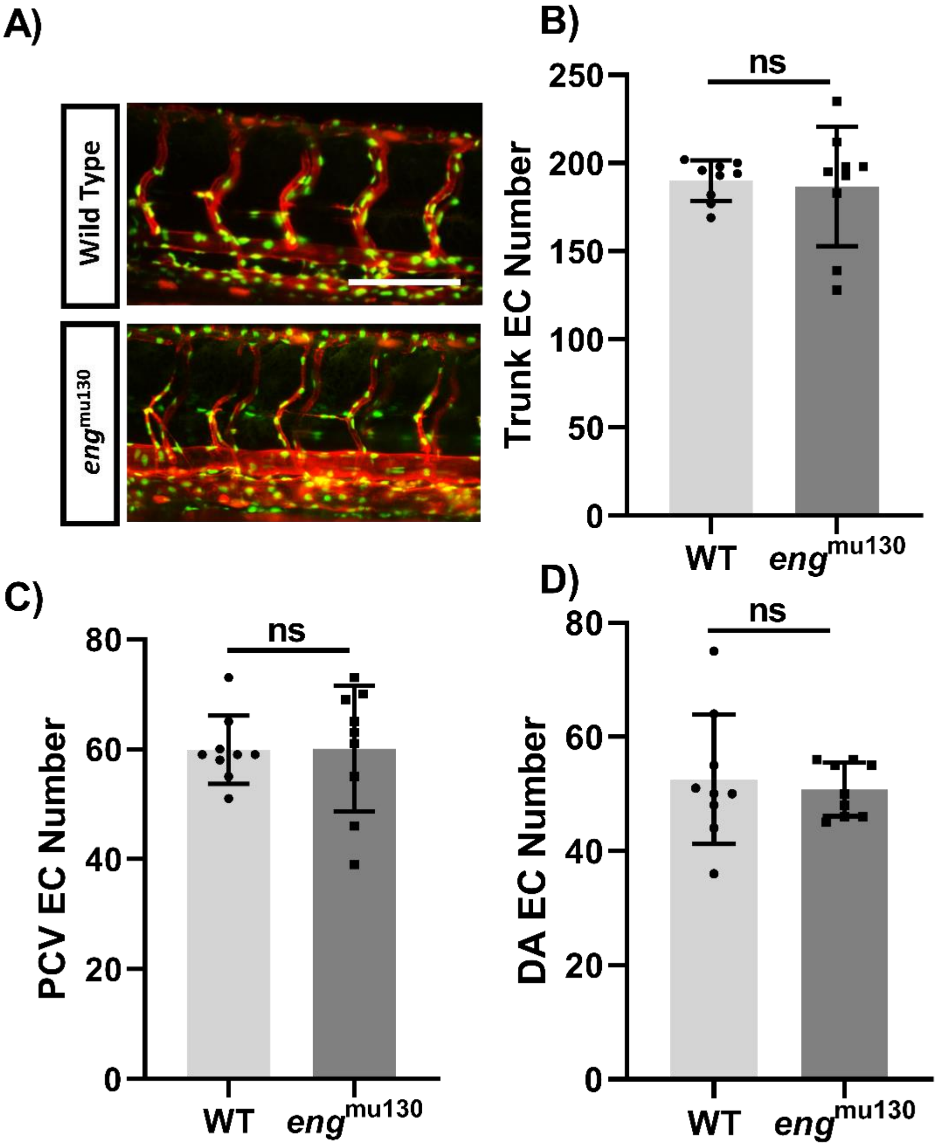


**Supplementary Figure 4 *endoglin* loss of function does not affect endothelial cell numbers.** A) Representative maximum intensity projection of *Tg(kdrl:Hsa.HRAS-mCherry)*$^{s916}$, Tg(fli1a:nEGFP)y7 trunk vasculature at 72hpf. B) Quantification of endothelial cell nuclei in all vessels in the region of interest. C) Quantification of endothelial cell nuclei in the dorsal aorta (DA). D) Quantification of endothelial cell nuclei in the posterior cardinal vein (PCV). (unpaired student's t-test. n =9/group.)

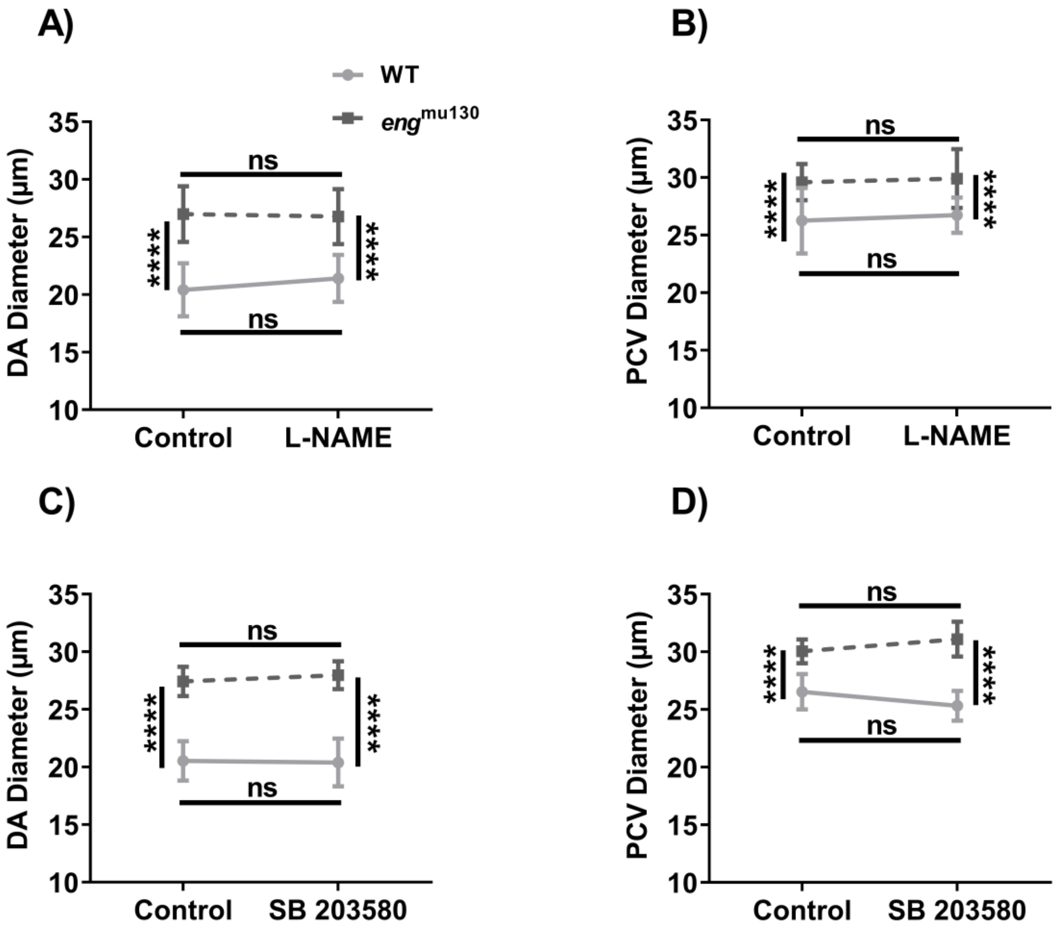


**Supplementary Figure 5 NOS or p38 MAPK inhibition between 2-3dpf does not affect the phenotype of *eng*$^{mu130}$ and WT embryos.** A B) Quantification of vessel diameter for the DA and PCV in WT and *eng*$^{mu130}$ mutant embryos with and without L-NAME treatment. C D) Quantification of vessel diameter for the DA and PCV in WT and *eng*$^{mu130}$ mutant embryos with and without SB 203580 treatment. (two-way ANOVA with Tukey post-hoc test. n = 10-15/group.)
